## Supporting Information for "Molecular principles of redox-coupled sodium pumping of the ancient Rnf machinery"

**Content**

**Extended Data Methods**
- Molecular simulations
- Electrostatic calculations
- Free energy profiles for $Na^+$ transport
- Simulation of the $Na^+$ translocation kinetics

**Extended Data Tables**
**Extended Data Table 1 |** Ferredoxin:$NAD^+$ oxidoreductase activities of purified Rnf-variants.
**Extended Data Table 2 |** Determination of the iron content for the WT Rnf complex and its variants.
**Extended Data Table 3 |** Growth rates and doubling time.
**Extended Data Table 4 |** Cryo-EM data collection, refinement, and validation statistics.
**Extended Data Table 5 |** Comparison of residues in NqrB of the Nqr complex and in RnfD of the Rnf complex.
**Extended Data Table 6 |** List of atomistic MD simulations.
**Extended Data Table 7 |** Non-standard protonation states in the MD simulations.
**Extended Data Table 8 |** Estimation of electron transfer rates.
**Extended Data Table 9 |** Plasmids generated during this study.

**Extended Data Figures**
**Extended Data Fig. 1 |** Purification and characterisation of the Rnf complex from *A. woodii*.
**Extended Data Fig. 2 |** Cryo-EM data collection and analysis of the Rnf complex with NADH bound.
**Extended Data Fig. 3 |** Cryo-EM data collection and analysis of the Rnf complex reduced with pre-reduced Fd.
**Extended Data Fig. 4 |** Cryo-EM data collection and analysis of the *apo* state of the Rnf complex.
**Extended Data Fig. 5 |** Cryo-EM density and model quality.
**Extended Data Fig. 6 |** Structural comparison of Rnf and Nqr and sequence conservation.
**Extended Data Fig. 7 |** MD simulations of the Rnf complex.
**Extended Data Fig. 8 |** Characterisation of the inward/outward conformations.
**Extended Data Fig. 9 |** Summary of distances between cofactors from MD simulations.
**Extended Data Fig. 10 |** Global dynamics of the Rnf complex from principal component analysis (PCA) of the MD simulations.
**Extended Data Fig. 11 |** Sodium binding in the membrane domain of Rnf.
**Extended Data Fig. 12 |** Kinetic simulations of the sodium translocation process.
**Extended Data Fig. 13 |** Purification and characterisation of Rnf variants from *A. woodii*.
**Extended Data Fig. 14 |** Sodium binding to the RnfA/E dimer.
**Extended Data Fig. 15 |** Overview of hydration in the membrane subunits of Rnf.
**Extended Data Fig. 16 |** Ion channel analysis in RnfA/E.

**Extended Data Movies**
**Movie 1 |** Representation of the segmented cryo-EM map of Rnf and its corresponding model.
**Movie 2 |** Sodium binding from the intracellular and extracellular sides.
**Movie 3 |** Inward/outward transition from MD simulations.

**Movie 4 |** Dominant normal modes from MD simulations of the NADH-reduced structures.
**Movie 5 |** Dominant normal modes from MD simulations of the Fd-reduced structures.

**Extended Data References**

### Extended Data Methods

#### Molecular simulations

Two different initial models were created starting from structures/cryo-EM density obtained from the NADH- or the ferredoxin (Fd)-reduced experimental conditions. For the NADH-reduced model, the resolved 3.3 Å cryo-EM structure (PDB ID: 9ERI) was embedded in a POPC membrane using CHARMM-GUI[47], and the system was hydrated in a TIP3P water box. $Na^+$ and $Cl^-$ ions were added to neutralise the system with an ion concentration of 250 mM. The Fd-reduced protein model was generated using molecular dynamics flexible fitting (MDFF)[48] by fitting the NADH-reduced structure to the density of the Fd-reduced Rnf using MDFF. To this end, the coordinates of the protein were relaxed to the cryo-EM map for 2 ns, with secondary structure restraints applied to protein residues. The protonation states of the protein sidechains and disulphide bridges were determined using PROPKA3 v. 3.4.0[40]. The FMN cofactors, located in RnfG and RnfD subunits were modelled by covalently linking them to $T185^{RnfG}$ and $T156^{RnfD}$ via a phosphodiester bond (Extended Data Fig. 1a). The full system comprised *ca.* 430,000 atoms. To probe the effect of cofactor redox state on protein conformation and sodium binding, the iron-sulphur clusters, FMN, or riboflavin cofactors were modelled in different redox states (see Extended Data Table 6). All simulations were performed using the CHARMM36 force field in combination with in-house DFT-based parameters for the cofactors[16,41,49]. All simulations were run using NAMD[43] versions 2.14 or 3.0 in an *NPT* ensemble with Nosé-Hoover-Langevin pressure control ($p$=1 atm) and Langevin thermostat ($T$=310 K), with the integration timestep set to 2 fs and rigid bonds set for hydrogens using the shakeH algorithm. Long-range electrostatic interactions were implemented using the Particle-Mesh Ewald (PME) method with the grid spacing set to 1 Å, while the switching and cutoff distances for the Lennard-Jones potential were set to 10 and 12 Å, respectively. The MD trajectories were analysed using VMD[45] and MDAnalysis[46].

#### Electrostatic calculations

Sodium binding energies were estimated using a Molecular Mechanics Poisson-Boltzmann Surface Area (PBSA/MM) model as implemented in APBS[50,51]. In this regard, MD snapshots with sodium bound next to the AE1 FeS centre were selected, with a cutoff of 7 Å, from simulations S9-S12 (Extended Data Table 6). The protein subunits RnfA, RnfE, the AE1 FeS centre, and the bound sodium were included in the models for estimation of the interaction energies. The solvent was modelled with a dielectric constant of 78.4, the NaCl concentration to 250 mM, whilst the protein interior was modelled as polarizable dielectric medium with an $\varepsilon$=4. The interactions between protein residues and sodium were described by explicit atomic point charges and Lennard-Jones interactions.

### Free energy profiles for $Na^+$ transport

Free energy profiles for the redox-driven $Na^+$ pumping in Rnf were derived based on redox potentials for Fd (-450 mV), FeS (-320 mV), FMN (-280 mV), RFB (-230 mV), and $NAD^+$ (-320 mV). The $Na^+$ affinity for the reduced AE1 centre was -130 mV (-3.1 kcal $mol^{-1}$), obtained from PBSA/MM calculations, which was also used to tune the redox potential of the AE1 centre to -190 mV in the $Na^+$ bound state to fulfil microscopic reversibility/detailed balance. The effect of the SMF on the free energy levels were divided equally based on the relative location of the cofactors in the membrane plane, $(z/L)\Delta\psi$, with the membrane thickness $L$ (32 Å) and the SMF ($\Delta\psi$) set to 180 mV. This ansatz yielded shifts in the electron transfer reactions for B8 → AE1 (+0.375$L$, $\Delta\Delta G$=-67.5 mV); AE1 → $FMN^G$ (+0.375$L$, $\Delta\Delta G$=-67.5 mV); $FMN^G$ → $FMN^D$ (-0.125$L$, $\Delta\Delta G$=+22.5 mV); $FMN^D$→ RBF (-0.25$L$, $\Delta\Delta G$=+45 mV); and RBF→ C1 (-0.375$L$, $\Delta\Delta G$= +67.5 mV), with the positive (+) directions defined from the N-side → P-side. Moreover, transferring the $Na^+$ ion from the inside to AE1 (+0.375$L$) and from AE1 to the cytoplasmic side (+0.625$L$) increased the free energy of the respective states by +67.5 mV and +112.5 mV at a 180 mV SMF.

The barriers for the electron transfer reactions were computed based on electron transfer theory[52] using the Moser-Dutton model[53],

$$\log k = 13 - (1.2-0.8\rho)\,(r-3.6) - 3.1(\Delta G+\lambda)^2 / \lambda \qquad (1a)$$

and for reactions with $\Delta G > 0$[53],

$$\log k = 13 - (1.2-0.8\rho)\,(r-3.6) - 3.1(-\Delta G+\lambda)^2 / \lambda - \Delta G/0.06 \qquad (1b)$$

where $\rho$ is the protein packing density (here assumed 0.76[53,54]), $r$ is the edge-to-edges distance, $\Delta G$ is the driving force for the electron transfer reaction, and $\lambda$ is the reorganisation energy (0.7 eV[53]). The electron transfer rates were converted into activation free energies ($\Delta G^{\ddagger}$) using transition state theory,

$$k = \kappa\,(k_B T/h)\,\exp(-\Delta G^{\ddagger}/RT) \qquad (2)$$

with $\kappa$=1, and $k_B T/h \sim 6.45\ ps^{-1}$. The rate of sodium binding to the reduced active AE1 site was estimated based on MD simulations to ca. 0.5 µs ($\Delta G^{\ddagger}$~0.4 eV)

### Simulation of the $Na^+$ translocation kinetics

Kinetic simulations were performed by numerical integration of the master-equation,

$$\frac{dp_i}{dt} = \sum_j k_{ji} p_j - \sum_j k_{ij} p_i \qquad (3)$$

for all transitions between energy levels $i$ and $j$ with forward ($k_{ij}$) and backward ($k_{ij}$) rates derived from the free energy profiles (Fig. 5b). The initial/final electron transfer steps were modeled as irreversible. The kinetic simulations were performed using COPASI[55]. See Extended Data Fig. 12 for further details on the kinetic simulations.

**Extended Data Tables**

**Extended Data Table 1 | Ferredoxin:NAD$^+$ oxidoreductase activities of purified Rnf-variants.** Activities are the mean of three independent biological replicates, measured in triplicates ($n$ = 3).

| Variant | $Fd_{red}$:NAD$^+$ [U mg$^{-1}$] |
|---|---|
| **Rnf complementation** | 7.1 ± 1.1 |
| **RnfΔAE1** | - |
| **RnfΔB1** | - |
| **RnfD D249A/N123A** | - |
| **RnfG T185A/Y113A** | - |
| **RnfA Y105A** | 3.4 ± 0.7 |
| **RnfE R67A** | 0.7 ± 0.2 |
| **RnfE L103G** | 0.8 ± 0.2 |

**Extended Data Table 2 | Determination of the iron content for the WT Rnf complex and its variants.** The measurements are the mean of three independent biological replicates, measured in triplicates ($n$ = 3).

| Variant | Mol Fe/mol Rnf |
|---|---|
| **Rnf complementation** | 41.8 ± 1.5 |
| **RnfΔAE1** | 40.4 ± 1.1 |
| **RnfΔB1** | 38.4 ± 2.2 |
| **RnfD D249A/N123A** | 41.9 ± 2.0 |
| **RnfG T185A/Y113A** | 42.3 ± 1.7 |
| **RnfA Y105A** | 42.1 ± 2.3 |
| **RnfE R67A** | 42.2 ± 1.9 |
| **RnfE L103G** | 42.9 ± 2.3 |

**Extended Data Table 3 | Growth rates and doubling time.** Data was obtained after growth on $H_2 + CO_2$ for ~ 90 h in complex medium containing 20 mM of NaCl. Values are the mean of three independent biological replicates, measured in triplicates ($n$ = 3).

| Variant | final $OD_{600}$ | Growth rate µ [h$^{-1}$] | Doubling time [h] |
|---|---|---|---|
| ***ΔpyrE*** | 0.477 ± 0.05 | 0.02 | 28 ± 1.7 |
| ***Δrnf*** | n.g.[1] | n.g.[1] | n.g.[1] |
| **Rnf complementation** | 0.453 ± 0.02 | 0.03 | 30 ± 2.0 |
| **RnfΔAE1** | n.g.[1] | n.g.[1] | n.g.[1] |
| **RnfΔB1** | n.g.[1] | n.g.[1] | n.g.[1] |
| **RnfD D249A/N123A** | n.g.[1] | n.g.[1] | n.g.[1] |
| **RnfG T185A/Y113A** | n.g.[1] | n.g.[1] | n.g.[1] |
| **RnfA Q85A** | 0.276 ± 0.02 | 0.015 | 43.5 ± 2.0 |
| **RnfA Y105A** | 0.175 ± 0.03 | 0.01 | 58.5 ± 2.5 |
| **RnfA T110G** | n.g.[1] | n.g.[1] | n.g.[1] |
| **RnfA T111G** | 0.411 ± 0.04 | 0.03 | 31.2 ± 1.3 |
| **RnfE R67A** | n.g.[1] | n.g.[1] | n.g.[1] |
| **RnfE L103G** | n.g.[1] | n.g.[1] | n.g.[1] |
| **RnfE V106G** | n.g.[1] | n.g.[1] | n.g.[1] |
| **RnfE N107A** | 0.150 ± 0.05 | 0.007 | 75.2 ± 3.1 |
| **RnfE E115A** | 0.390 ± 0.04 | 0.03 | 34.4 ± 2.2 |
| **RnfE E115Q** | 0.200 ± 0.05 | 0.01 | 65.3 ± 3.9 |
| **RnfE E115K** | n.g.[1] | n.g.[1] | n.g.[1] |
| **RnfD I274A** | 0.479 ± 0.07 | 0.03 | 31.2 ± 3.0 |
| **RnfD R275A** | 0.449 ± 0.03 | 0.02 | 30.7 ± 2.8 |
| **RnfD Y280A** | 0.440 ± 0.02 | 0.03 | 33.1 ± 4.0 |
| **RnfD L241A** | 0.457 ± 0.02 | 0.03 | 32.4 ± 3.5 |
| **RnfD F245A** | 0.455 ± 0.03 | 0.02 | 29.5 ± 3.0 |

[1]n. g., no growth was observed

**Extended Data Table 4 | Cryo-EM data collection, refinement, and validation statistics.**

| | **Rnf NADH bound state** (EMDB - 19915) (PDB - 9ERI) | **Rnf Fd-reduced State-1 (consensus map)** (EMDB - 19919) (PDB - 9ERK) | **Rnf Fd-reduced State-2 (B8 closer to membrane)** (EMDB - 19916) (PDB - 9ERJ) | **Rnf *apo* state** (EMDB - 19920) (PDB - 9ERL) |
|---|---|---|---|---|
| **Data collection and processing** | | | | |
| Magnification | 60,000 x | 165,000 x | 165,000 x | 165,000 x |
| Voltage (kV) | 300 | 300 | 300 | 300 |
| Electron exposure (e–/Å$^2$) | 50.0 | 50.0 | 50.0 | 50.0 |
| Defocus range (μm) | 0.8-1.8 | 1.2-1.8 | 1.2-1.8 | 1.2-1.8 |
| Pixel size (Å) | 1.09 | 0.75 | 0.75 | 0.84 |
| Symmetry imposed | C1 | C1 | C1 | C1 |
| Initial particle images (no.) | 3622306 | 5365845 | 5365845 | 2513935 |
| Final particle images (no.) | 645102 | 604534 | 260238 | 251476 |
| Map resolution (Å) | 3.3 | 2.8 | 2.99 | 3.0 |
| FSC threshold | 0.143 | 0.143 | 0.143 | 0.143 |
| Map resolution range (Å) | 3.0-4.5 | 2.5-4.0 | 2.8-4.0 | 2.8-4.0 |
| **Refinement** | | | | |
| Initial model used (PDB code) | *de novo,* AlphaFold | *de novo*, AlphaFold | *de novo*, AlphaFold | *de novo*, AlphaFold |
| Model resolution (Å) | 3.4 | 3.0 | 3.2 | 3.2 |
| FSC threshold | 0.5 | 0.5 | 0.5 | 0.5 |
| Model resolution range (Å) | 3.3-3.6 | 2.8-3.0 | 2.9-3.2 | 3.2-3.6 |
| Map sharpening *B* factor (Å$^2$) | -209.8 | -132.6 | -110.9 | -112.7 |
| Model composition | | | | |
| Non-hydrogen atoms | 12706 | 12669 | 12665 | 12665 |
| Protein residues | 1688 | 1688 | 1688 | 1688 |
| Ligands | FMN: 3, NADH: 1, RBF: 1, FES: 1, SF4: 10 | FMN: 3, NADH: 0, RBF: 1, FES: 1, SF4: 10, Na: 7 | FMN: 3, NADH: 0, RBF: 1, FES: 1, SF4: 10, Na: 3 | FMN: 3, NADH: 0, RBF: 1, FES: 1, SF4: 10, Na: 3 |
| *B* factors (Å$^2$) | | | | |
| Protein | 47.73 | 31.78 | 38.81 | 12.80 |
| Ligand | 50.96 | 51.81 | 70.93 | 22.86 |
| R.m.s. deviations | | | | |
| Bond lengths (Å) | 0.005 | 0.007 | 0.007 | 0.011 |
| Bond angles (°) | 0.778 | 1.159 | 1.097 | 1.231 |
| Validation | | | | |
| MolProbity score | 1.72 | 1.64 | 1.62 | 1.61 |
| Clashscore | 5.40 | 3.62 | 3.47 | 4.01 |
| Poor rotamers (%) | 0.58 | 0.52 | 0.45 | 0.67 |
| Ramachandran plot | | | | |
| Favored (%) | 96.44 | 96.00 | 96.24 | 95.50 |
| Allowed (%) | 3.02 | 3.28 | 3.22 | 4.08 |
| Disallowed (%) | 0.54 | 0.72 | 0.54 | 0.42 |

**Extended Data Table 5 | Comparison of residues present in NqrB of the Nqr complex and in RnfD of the Rnf complex.** Residues in NqrB involved in sodium translocation and the corresponding residues in RnfD are indicated. Conserved residues in $Na^+$- and $H^+$ -dependent Rnfs are highlighted. Sequence alignments were done with ClustalOmega[56].

| **Residue in NqrB** | **Residue in RnfD** | **Conserved residue in $Na^+$- and $H^+$ -dependent Rnfs** |
|---|---|---|
| **Na-1** | | |
| A263 | T177 | -[1] |
| V275 | I189 | -[1] |
| V332 | V235 | -[1] |
| **Na-2** | | |
| I371 | I274 | yes |
| R372 | R275 | yes |
| P376 | -[1] | |
| Y378 | Y280 | yes |
| F338 | L241 | yes |
| F342 | F245 | yes |

[1]-: residue not present in the sequence or not conserved in $Na^+$- and $H^+$ -dependent Rnfs

**Extended Data Table 6 | List of atomistic MD simulations.** The cumulative MD sampling was 8.3 µs. [a] Model generated based on initial MDFF relaxation. [b] See Extended Data Fig. 9 for classification of conformation.

| **Simulation** | **PDB** | **Substrate** | **Reduced cofactor** | **RnfA/E conformation**[b] | **Length (ns)** |
|---|---|---|---|---|---|
| S0 | 9ERI (nadh) | $NAD^+$ | - | outward | 700 |
| S1 | 9ERI (nadh) | $NAD^+$ | B8 | outward | 500 |
| S2 | 9ERI (nadh) | $NAD^+$ | B8 | outward | 500 |
| S3 | 9ERI (nadh) | $NAD^+$ | AE1 | outward | 500 |
| S4 | 9ERI (nadh) | $NAD^+$ | AE1 | outward | 500 |
| S5 | 9ERI (nadh) | $NAD^+$ | AE1, B8 | outward | 500 |
| S6 | 9ERI (nadh) | $NAD^+$ | AE1, B8 | outward | 500 |
| S7 | 9ERJ (fd)[a] | - | B7, B8 | outward | 500 |
| S8 | 9ERJ (fd) | - | B7, B8 | inward | 500 |
| S9 | 9ERJ (fd) | - | B8, AE1 | inward | 500 |
| S10 | 9ERJ (fd) | - | B8, AE1 | outward | 500 |
| S11 | 9ERJ (fd) | - | AE1, $FMN^G$ | outward | 500 |
| S12 | 9ERJ (fd) | - | AE1, $FMN^G$ | inward | 500 |
| S13 | 9ERJ (fd) | - | $FMN^G$, $FMN^D$ | inward | 500 |
| S14 | 9ERJ (fd) | - | $FMN^G$, $FMN^D$ | inward | 500 |
| S15 | 9ERJ (fd) | - | B8, $FMN^G$ | inward | 100 |
| S16 | 9ERJ (fd) | - | B8, $FMN^G$ | outward | 100 |
| S17 | 9ERJ (fd) | - | B8, $FMN^D$ | inward | 100 |
| S18 | 9ERJ (fd) | - | B8, $FMN^D$ | inward | 100 |
| S19 | 9ERJ (fd) | - | B8, RBF | inward | 100 |
| S20 | 9ERJ (fd) | - | B8, RBF | inward | 100 |
| S21 | 9ERJ (fd) | - | RBF, $FMN^D$ | inward | 500 |
| S22 | 9ERJ (fd) | - | RBF, $FMN^D$ | outward | 500 |
| Total | | | | | 8300 ns |

**Extended Data Table 7 | Non-standard protonation states in the MD simulations.** The protonation states were determined using PROPKA3 calculation based on the NADH-reduced cryo-EM structure. (ε) - ε protonated (neutral) histidine, (δ) - δ protonated (neutral) histidine, (ε/δ) - positively charged doubled protonated histidine ($HisH^+$). Unless stated otherwise, histidine residues were modelled in δ-protonated state.

| **Subunit** | **Residues** |
|---|---|
| RnfA | - |
| RnfB | E127 |
| RnfC | E167, E169, H5(ε/δ) |
| RnfD | E61, E188, D249, H139(ε/δ) |
| RnfE | - |
| RnfG | E87, E139 |

**Extended Data Table 8 | Estimation of electron transfer rates.** Redox potentials: Fd (-450 mV), FeS (-320 mV), FMN (-280 mV), RFB (-230 mV), $NAD^+$ (-320 mV) [a] State is not sampled. <*r*> – average *edge-to-edge* distances from MD simulations; the $\lambda$ – reorganisation energy was modelled as 0.7 eV[53]; $\rho$ – protein packing density was set to 0.76[53]; $\Delta G^{\ddagger}$ – Activation free energies based on transition state theory from $k_{eT}$, with standard pre-exponential factors and $\kappa$=1; electron transfer rate $k_{eT}$ from Moser-Dutton model[53] (see also Ref. 60). [c] shortest distance from MD snapshots. [d] based on FMN/NADH PCET reaction in Complex I, ca. 10,000 $s^{-1}$ [17].

| **D** | **A** | **State** | **<*r* (Å)>** | **Δ*G* (eV)** | **$\Delta G^{\ddagger}$ (eV)** | **$t_{eT}$ (s)** | **Δ*G* (eV) SMF** | **$\Delta G^{\ddagger}$ (eV) SMF** | **$t_{eT}$ (s) SMF** |
|---|---|---|---|---|---|---|---|---|---|
| Fd | B1 | *N/A*[a] | 14[a] | -0.13 | 0.46 | $3.9\times10^{-6}$ | -0.13 | 0.46 | $3.9\times10^{-6}$ |
| B1 | B2 | *inward* | 9.6 | 0 | 0.34 | $5.3\times10^{-8}$ | 0 | 0.34 | $5.3\times10^{-8}$ |
| B1 | B2 | *outward* | 11.7 | 0 | 0.29 | $9.2\times10^{-9}$ | 0 | 0.29 | $9.2\times10^{-9}$ |
| B2 | B3 | *inward* | 13.0 | 0 | 0.28 | $5.4\times10^{-9}$ | 0 | 0.28 | $5.4\times10^{-9}$ |
| B2 | B3 | *outward* | 16.3 | 0 | 0.58 | $0.5\times10^{-3}$ | 0 | 0.58 | $0.5\times10^{-3}$ |
| B3 | B4 | *inward* | 11.1 | 0 | 0.40 | $4.1\times10^{-7}$ | 0 | 0.40 | $4.1\times10^{-7}$ |
| B3 | B4 | *outward* | 10.6 | 0 | 0.38 | $2.1\times10^{-7}$ | 0 | 0.38 | $2.1\times10^{-7}$ |
| B4 | B5 | *inward* | 11.3 | 0 | 0.40 | $5.3\times10^{-7}$ | 0 | 0.40 | $5.3\times10^{-7}$ |
| B4 | B5 | *outward* | 11.2 | 0 | 0.40 | $4.7\times10^{-7}$ | 0 | 0.40 | $4.7\times10^{-7}$ |
| B5 | B6 | *inward* | 10.5 | 0 | 0.37 | $1.8\times10^{-7}$ | 0 | 0.37 | $1.8\times10^{-7}$ |
| B5 | B6 | *outward* | 10.8 | 0 | 0.38 | $2.7\times10^{-7}$ | 0 | 0.38 | $2.7\times10^{-7}$ |
| B6 | B7 | *inward* | 10.9 | 0 | 0.39 | $3.1\times10^{-7}$ | 0 | 0.39 | $3.1\times10^{-7}$ |
| B6 | B7 | *outward* | 11.3 | 0 | 0.40 | $5.4\times10^{-7}$ | 0 | 0.40 | $5.4\times10^{-7}$ |
| B7 | B8 | *inward* | 11.0 | 0 | 0.39 | $3.6\times10^{-7}$ | 0 | 0.39 | $3.6\times10^{-7}$ |
| B7 | B8 | *outward* | 9.1 | 0 | 0.32 | $2.7\times10^{-8}$ | 0 | 0.32 | $2.7\times10^{-8}$ |
| B8 | AE1 | *inward* | 15[c] | 0 | 0.54 | $8.3\times10^{-5}$ | -0.07 | 0.51 | $3.2\times10^{-5}$ |
| B8 | AE1 | *inward* + $Na^+$ | 15[c] | -0.26 | 0.46 | $4.0\times10^{-6}$ | -0.13 | 0.49 | $1.5\times10^{-5}$ |
| B8 | AE1 | *outward* | 20.7 | 0 | 0.74 | 0.2 | -0.07 | 0.72 | 0.08 |
| B8 | AE1 | *outward* + $Na^+$ | 20.7 | -0.26 | 0.66 | 0.01 | -0.13 | 0.70 | 0.04 |
| AE1 | $FMN^G$ | *inward* | 23.1 | +0.09 | 0.87 | 20.3 | +0.13 | 0.89 | 39.4 |
| AE1 | $FMN^G$ | *outward* | 15.1 | +0.09 | 0.58 | $0.4\times10^{-3}$ | +0.13 | 0.59 | $0.7\times10^{-3}$ |
| $FMN^G$ | $FMN^D$ | inward | 11.6 | 0 | 0.41 | $8.1\times10^{-7}$ | +0.03 | 0.42 | $1.2\times10^{-6}$ |
| $FMN^G$ | $FMN^D$ | outward | 12.1 | 0 | 0.43 | $1.6\times10^{-6}$ | +0.03 | 0.44 | $2.5\times10^{-6}$ |
| $FMN^D$ | RBF | inward | 10.6 | -0.05 | 0.36 | $1.0\times10^{-7}$ | 0 | 0.38 | $2.1\times10^{-7}$ |
| $FMN^D$ | RBF | outward | 10.9 | -0.05 | 0.37 | $1.6\times10^{-7}$ | 0 | 0.39 | $3.1\times10^{-7}$ |
| RBF | C1 | *inward* | 11.3 | +0.09 | 0.44 | $2.1\times10^{-6}$ | +0.1 | 0.44 | $2.5\times10^{-6}$ |
| RBF | C1 | *outward* | 11.4 | +0.09 | 0.44 | $2.4\times10^{-6}$ | +0.1 | 0.45 | $2.8\times10^{-6}$ |
| C1 | C2 | *inward* | 9.4 | 0 | 0.33 | $4.0\times10^{-8}$ | 0 | 0.33 | $4.0\times10^{-8}$ |
| C1 | C2 | *outward* | 9.6 | 0 | 0.34 | $5.3\times10^{-8}$ | 0 | 0.34 | $5.3\times10^{-8}$ |
| C1 | FMN | *inward* | 8.5 | +0.04 | 0.32 | $2.1\times10^{-8}$ | +0.04 | 0.32 | $2.1\times10^{-8}$ |
| C1 | FMN | *outward* | 8.5 | +0.04 | 0.32 | $2.1\times10^{-8}$ | +0.04 | 0.32 | $2.1\times10^{-8}$ |
| FMN | $NAD^+$ | *inward* | **3.5** | -0.04 | 0.54 | $0.1\times10^{-3}$ | -0.04 | 0.54 | $0.1\times10^{-3}$ |
| FMN | $NAD^+$ | *outward* | **3.5** | -0.04 | 0.54 | $0.1\times10^{-3}$ | -0.04 | 0.54 | $0.1\times10^{-3}$ |

**Extended Data Table 9 | Plasmids generated during this study.**

| Plasmid | Description | Number of oligonucleotides used[1] | Introduced mutations |
|---|---|---|---|
| pMTL84211_pPta_ack_ RnfCDGEAB (RnfG-His) | WT Rnf, C-terminal His-tag fused to RnfG | 1+2+3+4+5+6 | None |
| pMTL84211_pPta_ack_Rnf ΔAE1 | Deletion of FeS cluster AE1 | 7+8+9+10+ 11+12+13+14 | A-C25A, A-C113A, E-C25A, E-C108A |
| pMTL84211_pPta_ack_Rnf ΔB1 | Deletion of FeS cluster B1 | 15+16 | C50A, C53A, C58A, C75A |
| pMTL84211_pPta_ack_Rnf D D249A/N123A | Exchange of Riboflavin-binding D249 and N123 to A249 and A123 in RnfD | 17+18+19+20 | D249A, N123A |
| pMTL84211_pPta_ack_ RnfG T185A/Y113A | Exchange of FMN-binding T185 and Y113 to A185 and A113 in RnfG | 21+22+23+24 | T185A, Y113A |
| pMTL84211_pPta_ack_Rnf A Y105A | Exchange of Y105 to A in RnfA | 25+26 | Y105A |
| pMTL84211_pPta_ack_Rnf A T110G | Exchange of T110 to G in RnfA | 27+28 | T110G |
| pMTL84211_pPta_ack_Rnf A T111G | Exchange of T111 to G in RnfA | 29+30 | T111G |
| pMTL84211_pPta_ack_Rnf A Q85A | Exchange of Q85 to A in RnfA | 31+32 | Q85A |
| pMTL84211_pPta_ack_Rnf E R67A | Exchange of R67 to A in RnfE | 33+34 | R67A |
| pMTL84211_pPta_ack_Rnf E L103G | Exchange of L103 to G in RnfE | 35+36 | L103G |
| pMTL84211_pPta_ack_Rnf E V106G | Exchange of V106 to G in RnfE | 37+38 | V106G |
| pMTL84211_pPta_ack_Rnf E N107A | Exchange of N107 to A in RnfE | 39+40 | N107A |
| pMTL84211_pPta_ack_Rnf E E115A | Exchange of E115 to A in RnfE | 41+42 | E115A |
| pMTL84211_pPta_ack_Rnf E E115Q | Exchange of E115 to Q in RnfE | 43+44 | E115Q |
| pMTL84211_pPta_ack_Rnf E E115K | Exchange of E115 to K in RnfE | 45+46 | E115K |
| pMTL84211_pPta_ack_Rnf D I274A | Exchange of I274 to A in RnfD | 47+48 | I274A |
| pMTL84211_pPta_ack_Rnf D R275A | Exchange of R275 to A in RnfD | 49+50 | R275A |
| pMTL84211_pPta_ack_Rnf D Y280A | Exchange of Y280 to A in RnfD | 51+52 | Y280A |
| pMTL84211_pPta_ack_Rnf D L241A | Exchange of L241 to A in RnfD | 53+54 | L241A |
| pMTL84211_pPta_ack_Rnf D F245A | Exchange of F245 to A in RnfD | 55+56 | F245A |

**Extended Data Figures**

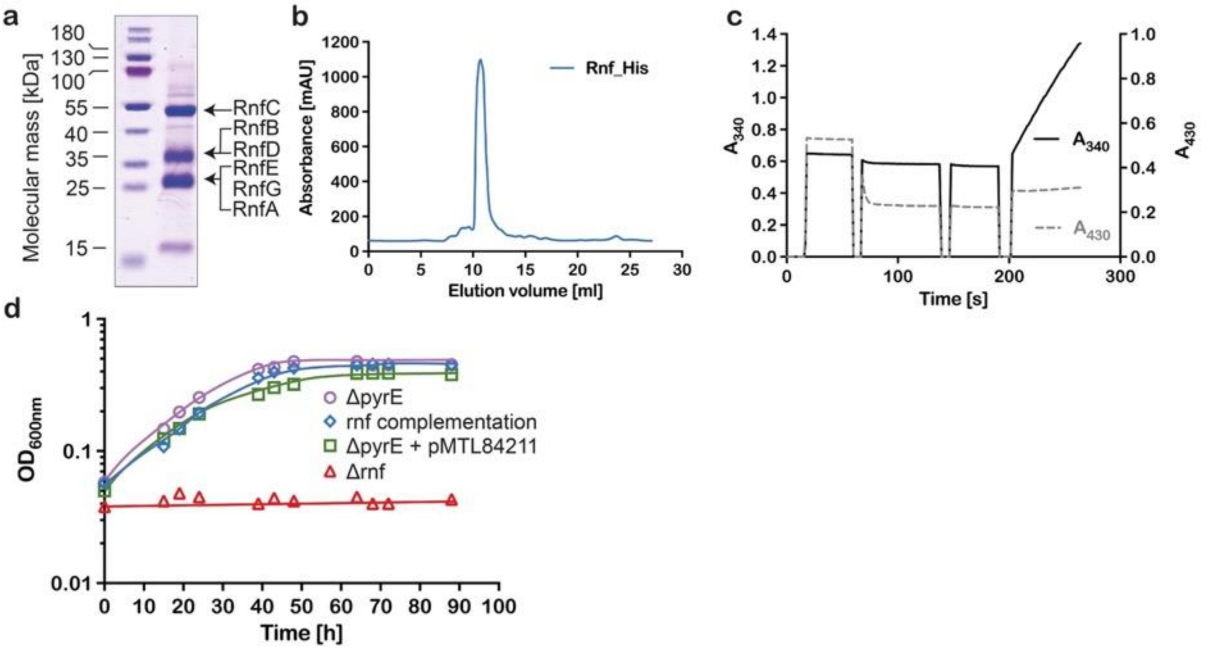

**Extended Data Fig. 1 | Purification and characterisation of the Rnf complex from *A. woodii*. (a)** 10 µg of Rnf complex containing a His-tag purified from *A. woodii* was separated in an SDS-PAGE. **(b)** Size exclusion chromatography profile of the purified Rnf complex from *A. woodii* on "Superdex 200 Increase™ 10/300". **(c)** Measurement of $Fd_{red}$-dependent $NAD^+$ reduction catalysed by the Rnf complex from *A. woodii*. **(d)** Growth restoration of the *rnf* mutant on $H_2$ and $CO_2$ after complementation with the plasmid pMTL84211_Ppta_ack_Rnf-His. The *A. woodii* Δ*pyrE* mutant, the Δ*pyrE* mutant containing the vector pMTL84211 without the *rnf* operon, the Δ*rnf* mutant and the complemented strain Δ*rnf* were grown in complex medium under a $H_2$ + $CO_2$ [80:20 v/v] atmosphere with a pressure of $1.0 \times 10^5$ Pa.

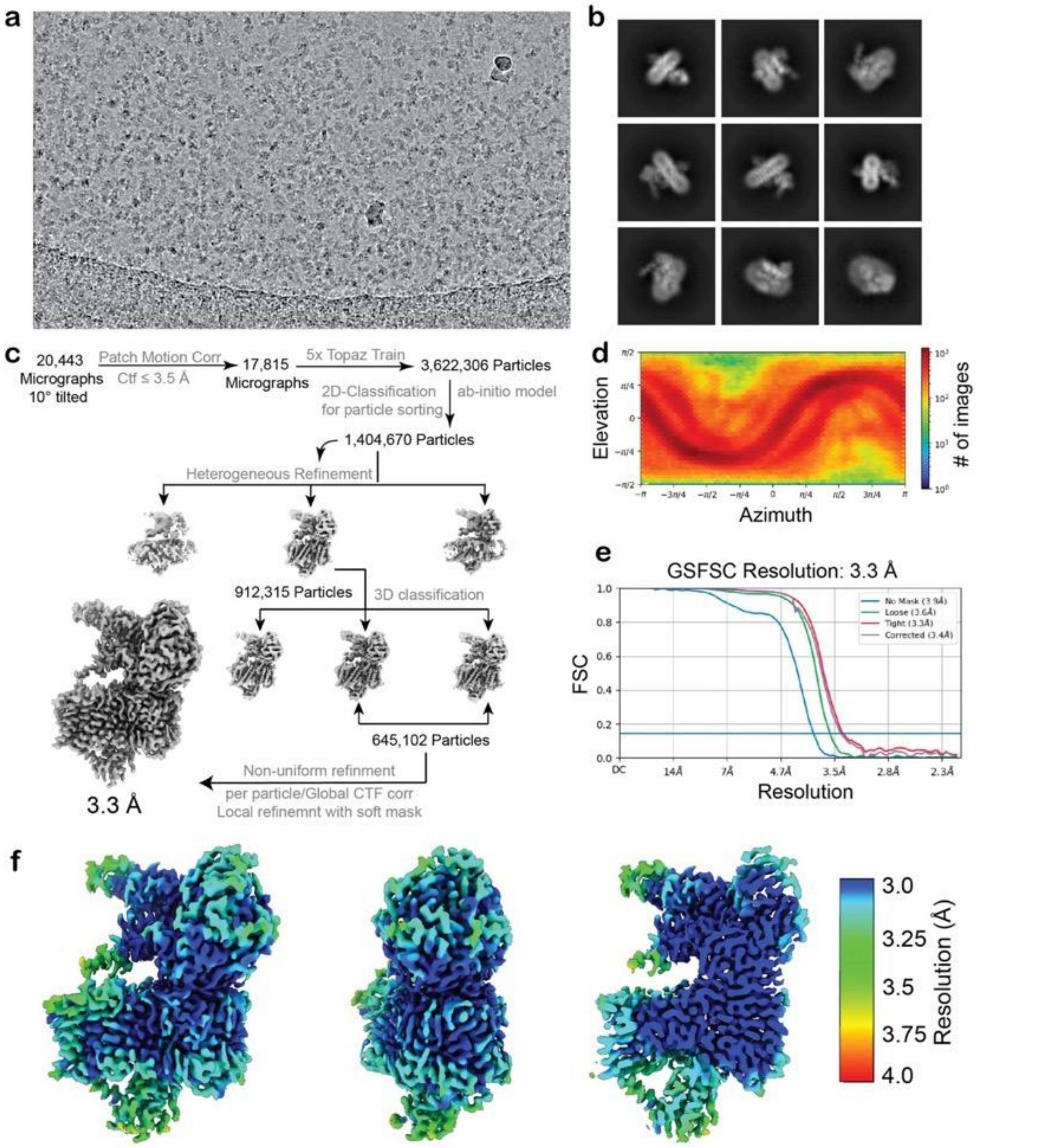

**Extended Data Fig. 2 | Cryo-EM data collection and analysis of the Rnf complex with NADH bound. (a)** A representative cryo-EM motion-corrected micrograph showing Rnf particles. **(b)** Reference-free 2D class averages revealing different views of the Rnf complex. **(c)** Overview of the cryo-EM data-processing scheme. A 10° tilted dataset was acquired to overcome the preferred orientation problem. **(d)** Angular distribution of the particles used for the final round of NU refinement. **(e)** Fourier shell correlation plot of final refined map with showing the global resolution (FSC = 0.143). **(f)** Local resolution as calculated by CryoSparc[32] mapped on the refined density (left, middle: front and side view, right: cut-open view of central section).

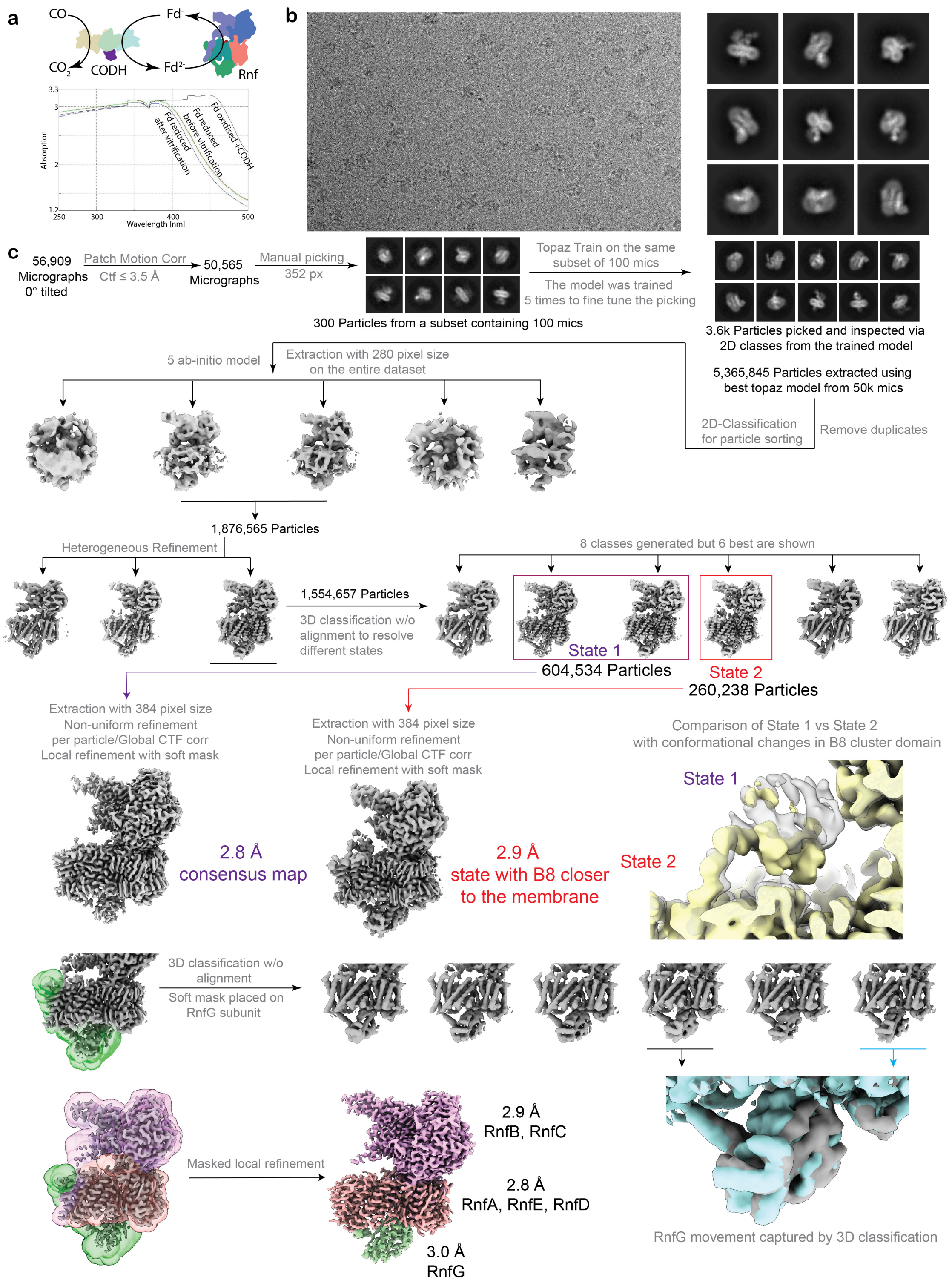

**Extended Data Fig. 3 | Cryo-EM data collection and analysis of the Rnf complex reduced with pre-reduced Fd. (a)** Cartoon scheme depicting the CODH enzyme extracting electrons from CO gas and transferring these electrons to oxidised Fd to produce reduced Fd. The Rnf is then incubated with reduced Fd to obtain a reduced protein complex. **(b)** A representative cryo-EM motion-corrected micrograph showing single Rnf complexes and reference-free 2D class averages revealing different views of the complex. **(c)** Overview of the cryo-EM data-processing scheme. Due to low number of particles on the grid, a large dataset was acquired to explore conformation changes. 3D classification revealed two conformations associated with the RnfB mobile domain containing the B8 cluster; one closer to the membrane (state 2) and one away (state 1) (*see methods* for a detailed description of the processing pipeline).

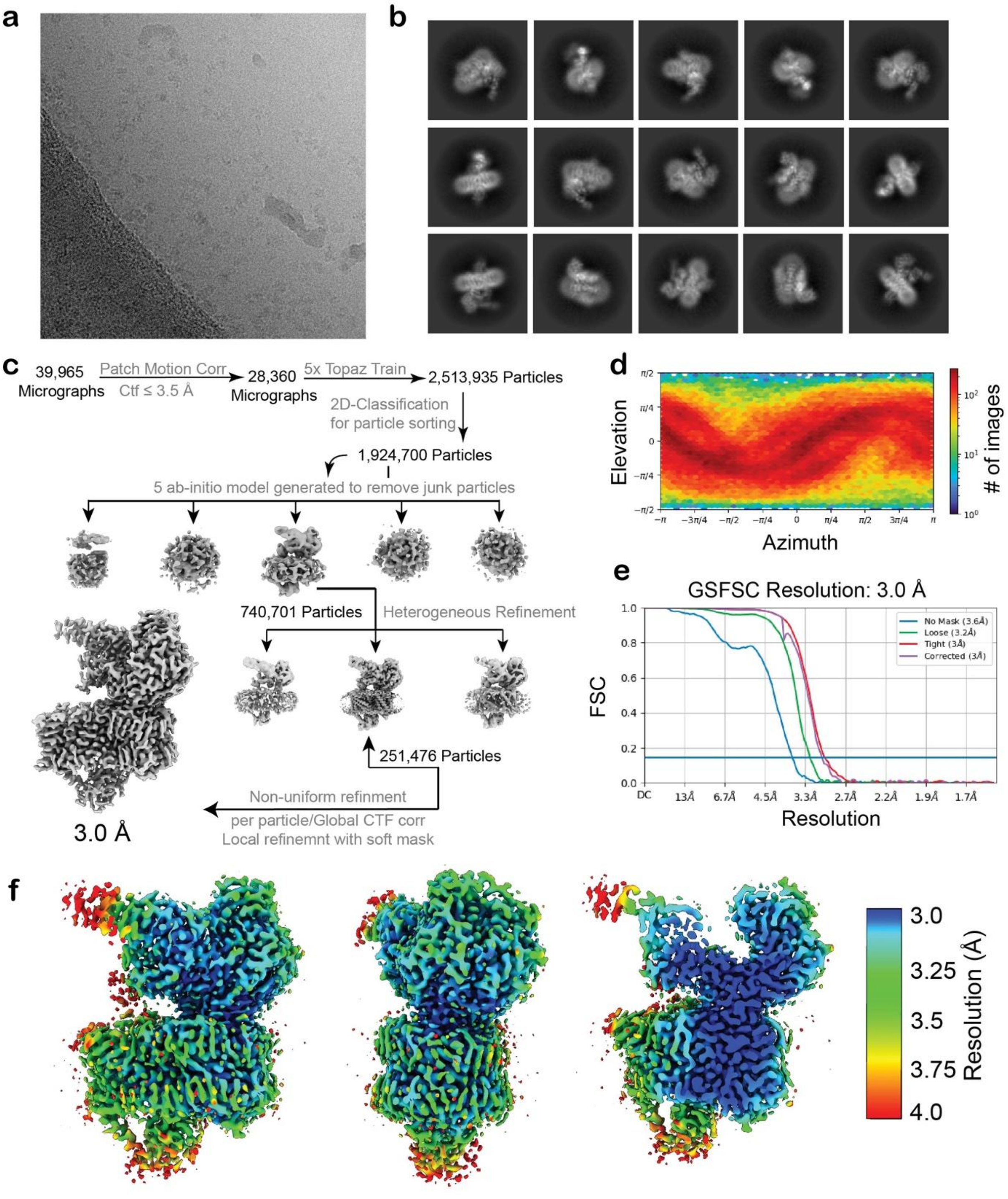

**Extended Data Fig. 4 | Cryo-EM data collection and analysis of the *apo* state of the Rnf complex. (a)** A representative cryo-EM micrograph showing Rnf particles. **(b)** Reference-free 2D class averages revealing different views of the Rnf. **(c)** Overview of the cryo-EM data-processing scheme. **(d)** Angular distribution of the particles used for the final round of NU refinement. **(e)** Fourier shell correlation plot of final refined map with showing the global resolution (FSC = 0.143). **(f)** Local resolution as calculated by CryoSparc[32] mapped on the refined density (*left*, middle: front and side view, *right*: cut-open view of central section).

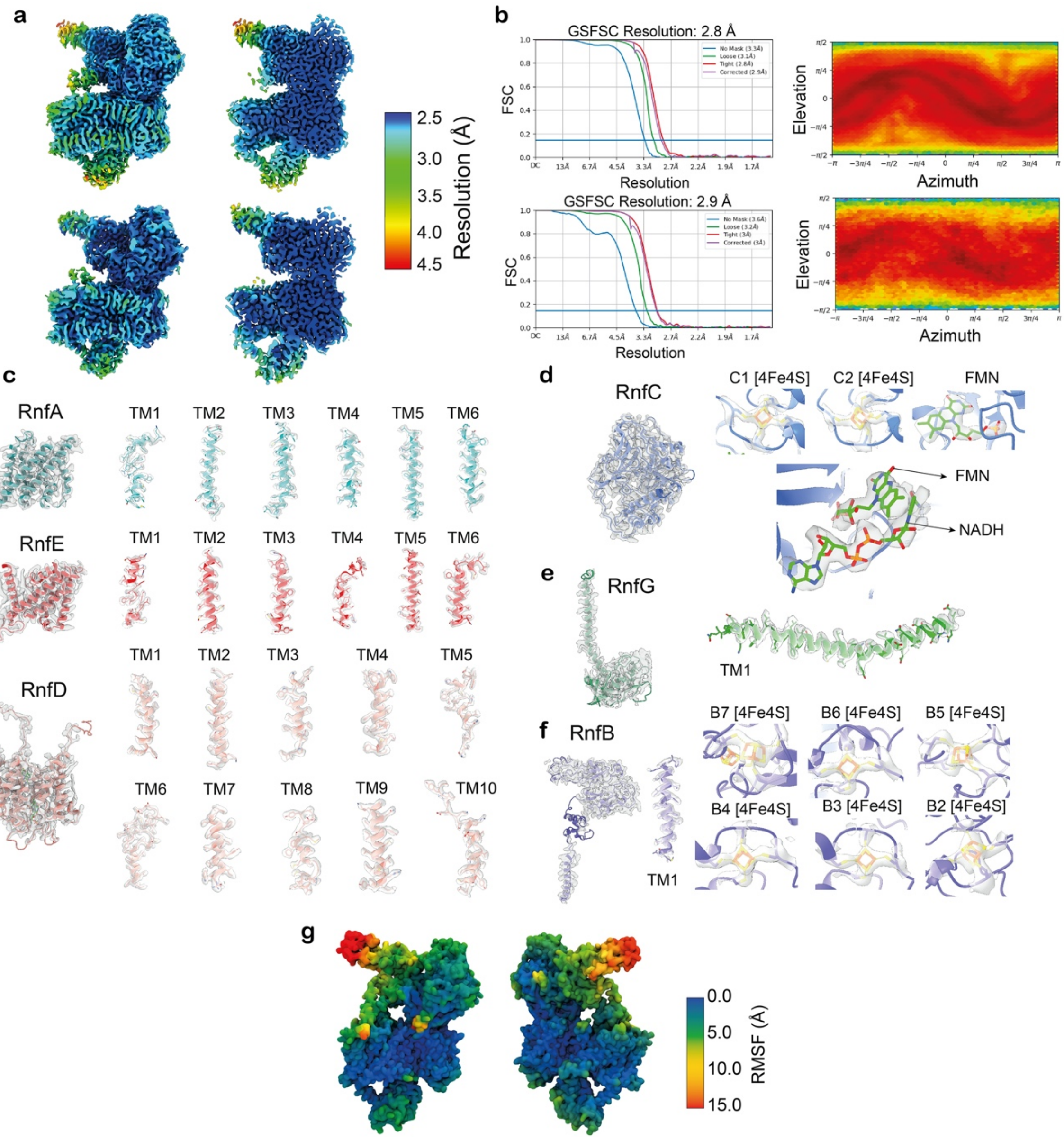

**Extended Data Fig. 5 | Cryo-EM density and model quality.** Representative regions of the Rnf subunits and their surrounding electron density maps are shown. **(a)** Local resolution as calculated by CryoSparc[32] mapped on the refined density (2.8 Å consensus map) of Rnf reduced with $Fd_{red}$ taken from Fig. S3 (front and cut-open view of central section). **(b, c, d, e, f)** Representative regions of the Rnf subunits and their surrounding electron density maps are shown. The helices TM 1 and 4 for RnfAE subunit were found to slightly disordered with moderate density fits, indicating their flexible nature. The FMN-NADH bound state was only obtained for Rnf complex with bound NADH, whereas for the Rnf in the Fd reduced state and the *apo* state contain only FMN. **(g)** Average root-mean-square fluctuations (RMSF) calculated from the MD simulations and mapped on the cryo-EM structure of the Rnf complex treated with reduced Fd.

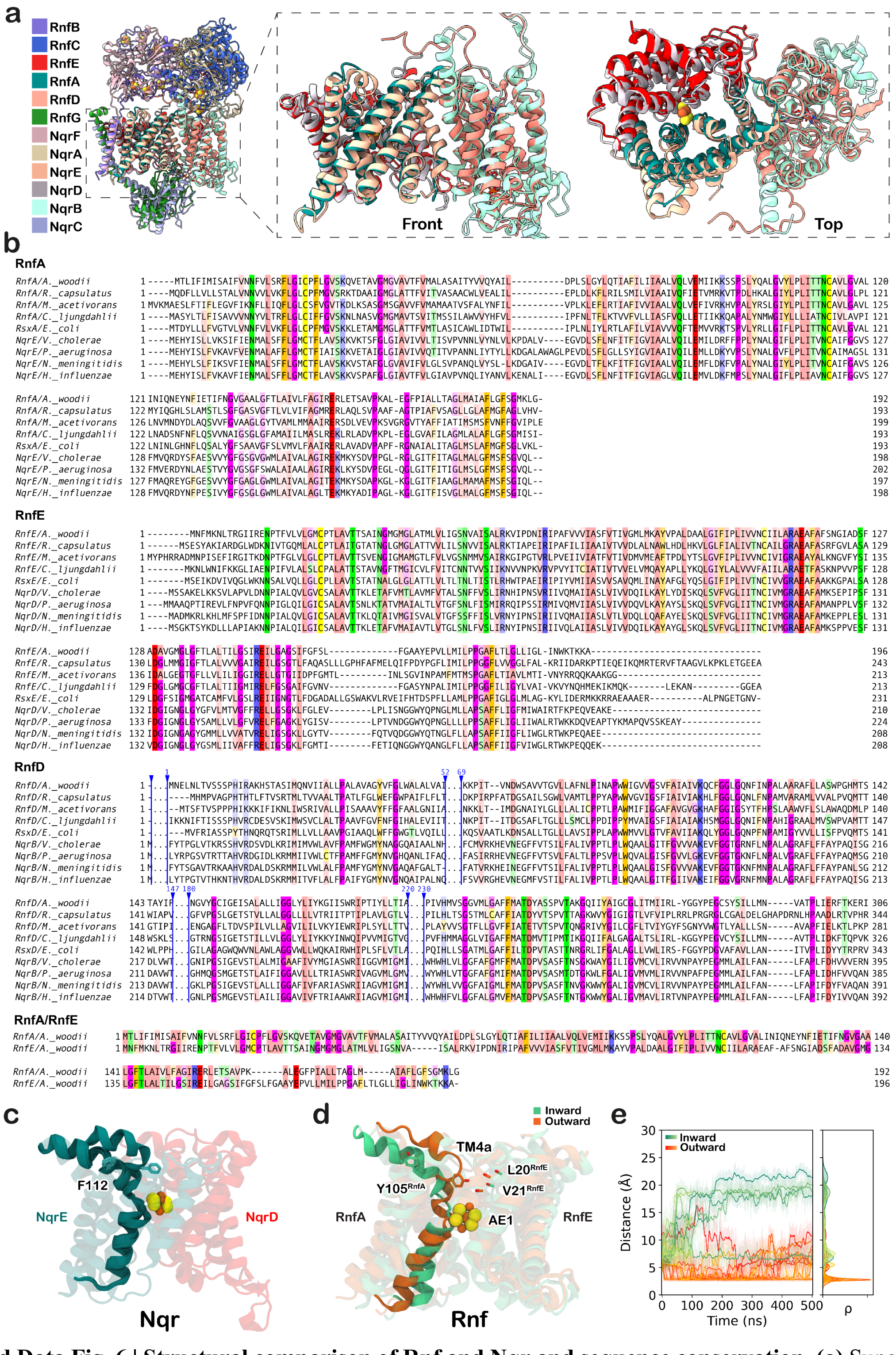

**Extended Data Fig. 6 | Structural comparison of Rnf and Nqr and sequence conservation. (a)** Superposition of the Rnf structure (PDB ID: 9ERI) with Nqr complex (PDB ID: 7XK3)[12]. The overall complex aligns with an RMSD of 4.75 Å, however, the membrane integral subunits, shown as closeup, are highly similar and align with an RMSD value of 1.18 Å. **(b)** The corresponding multiple-sequence alignment (MSA) of the membrane integral subunit of Rnf complex (RnfA/E/D) and Nqr complex (NqrB/D/E). The RnfA/E, although being pseudo-symmetrical, show a sequence similarity of 29%. The MSA was performed with ClustalΩ[56]. Residues with up to 50% conservation are coloured. **(c, d, e)** Comparison of conformations from resolved cryo-EM structures of Nqr (PDB ID: 8A1T, 8A1U, 8A1V, 8A1W, 8A1X, 8A1Y, 8ACW, 8ACY)[13] (c) with MD *inward/outward* conformations (d), (e) Distance between the residues Y105$^{RnfA}$-V21$^{RnfE}$ from MD simulations grouped either into *inward* (green) or *outward* (orange) conformations.

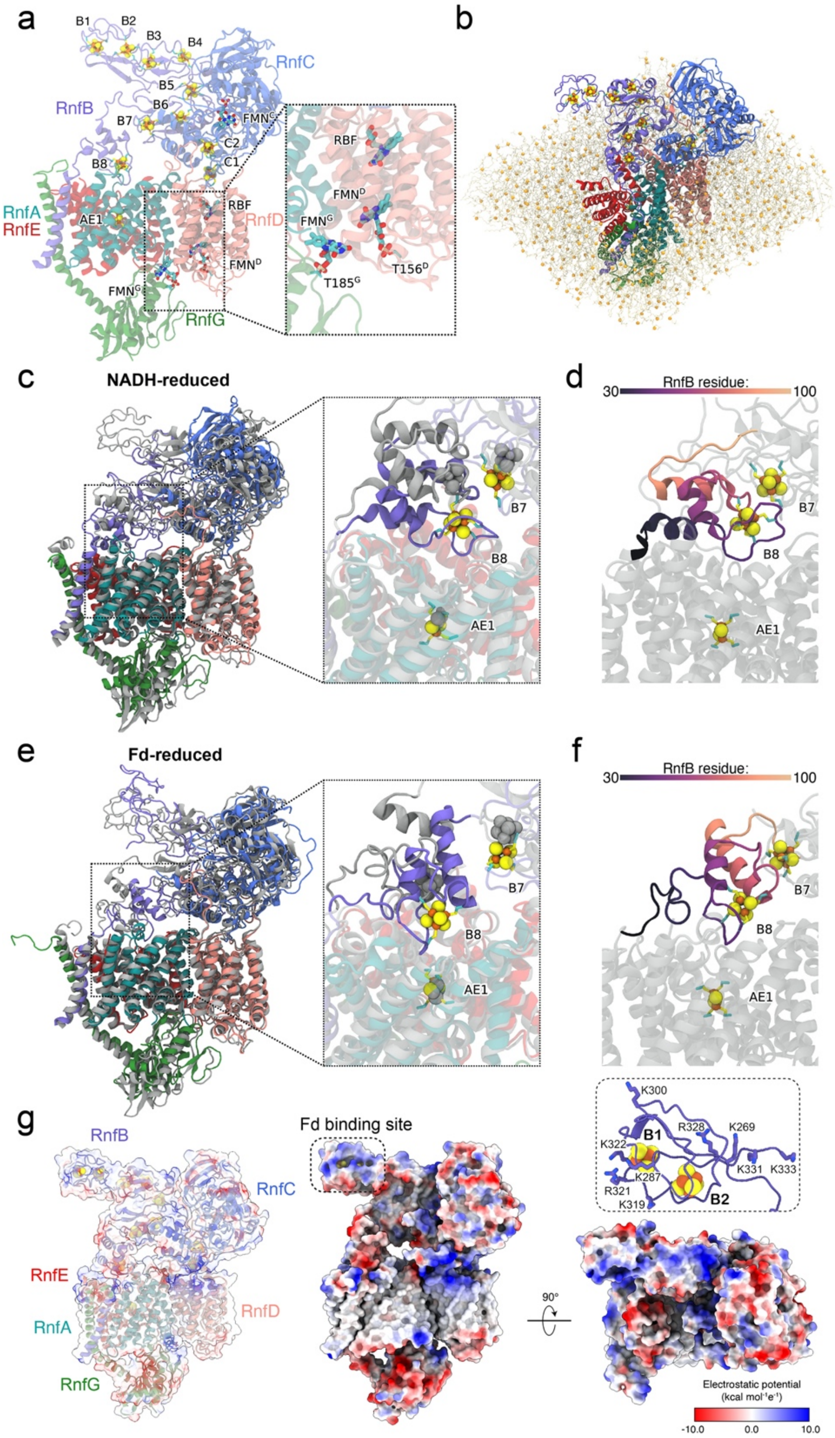

**Extended Data Fig. 7 | MD simulations of the Rnf complex. (a)** Structural overview of the Rnf system used for MD simulations. Subunits RnfA (teal), RnfB (purple), RnfC (blue), RnfD (pink), RnfE (red), RnfG (green) are shown in cartoon representation. The ISCs are represented as spheres (sulphur:yellow, iron:orange), while the cofactors FMN and RBF are shown as sticks. *Inset*: The $FMN^G$ and $FMN^D$ were covalently linked to $T185^G$ and $T156^D$, respectively. **(b)** The protein system embedded in a POPC membrane. Waters and ions are omitted for clarity. **(c,e)** Comparison of structures from cryo-EM experiment (grey) and equilibrated MD simulations. The MD simulations were started either from **(c)** the NADH- or **(e)** the Fd-reduced cryo-EM structures. *Inset:* close-up of B8 cluster binding region of RnfB (purple) showing the shift in position from MD simulations as compared to initial cryo-EM structure (grey). **(d, f)** Secondary structure of RnfB region binding the B8 cluster from MD simulations started from **(d)** the NADH- or **(f)** the Fd-reduced structures. Residue range of 30 (dark purple) to 100 (yellow) is shown. **(g)** Electrostatic potential on the Rnf surface and RMSF. RnfB shows a positive electrostatic potential region that could stabilise ferredoxin interaction. *Inset:* The C-terminal domain of RnfB contains several positive Lys/Arg residues.

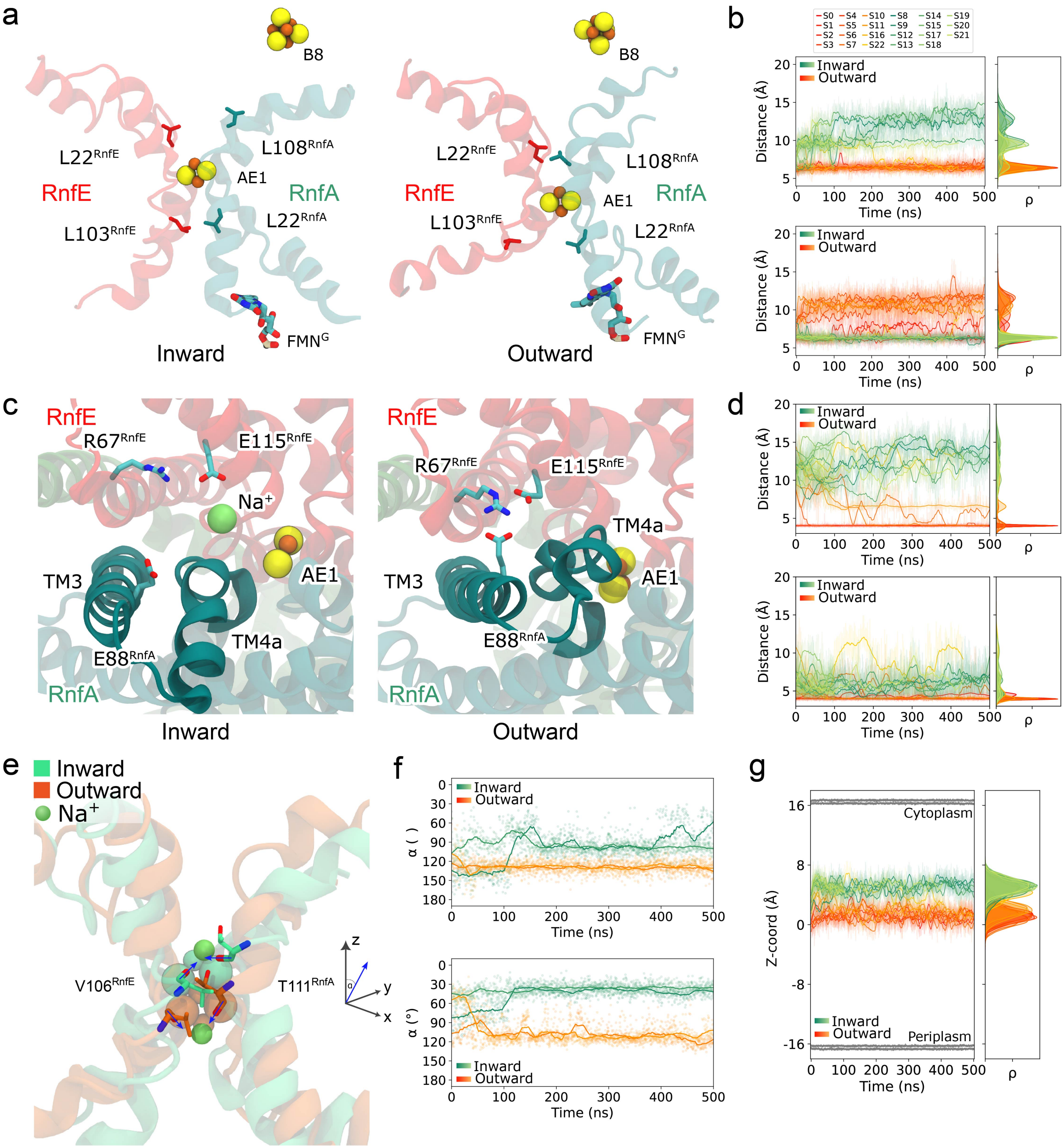

**Extended Data Fig. 8 | Characterisation of the inward and outward conformations. (a)** *Inward* (*left*) and *outward* (*right*) conformations of TM4 and TM1 helices from RnfA (teal) and RnfE (red). Conserved hydrophobic residues ($L22^{RnfA}$, $L22^{RnfE}$, $L103^{RnfE}$, $L108^{RnfA}$) can adopt different conformations depending on the helix conformation, controlling access to the AE1 cluster. **(b)** Distance between the gating hydrophobic residues *top:* $L22^{RnfE}$ and $L108^{RnfA}$; and *bottom*: $L22^{RnfA}$ and $V106^{RnfE}$. **(c)** Snapshots from the *inward* (left) and *outward* (right) conformations showing different ion pair conformations for $R67^{RnfE}$, $E88^{RnfA}$, and $E115^{RnfE}$. **(d)** Distances between the $R67^{RnfE}$-$E88^{RnfA}$ (top) and $R67^{RnfE}$-$E115^{RnfE}$ (bottom) ion pairs. **(e)** Sodium binding into the buried binding site is stabilised by backbone carbonyls $V106^{RnfE}$ and $T111^{RnfA}$ in the inward (green) and outward (orange) conformations. **(f)** Angle of backbone carbonyl from $V106^{RnfE}$ (top) and $T111^{RnfA}$ (bottom) projected onto a Z-axis (in Å) from MD simulations with a reduced AE1 cluster (S9-S12, see Extended Data Table 6). **(g)** The Z-coordinate (in Å) of the AE1 cluster relative to the membrane. The average Z-coordinate (in Å) of the upper and lower membrane leaflets are shown in grey.

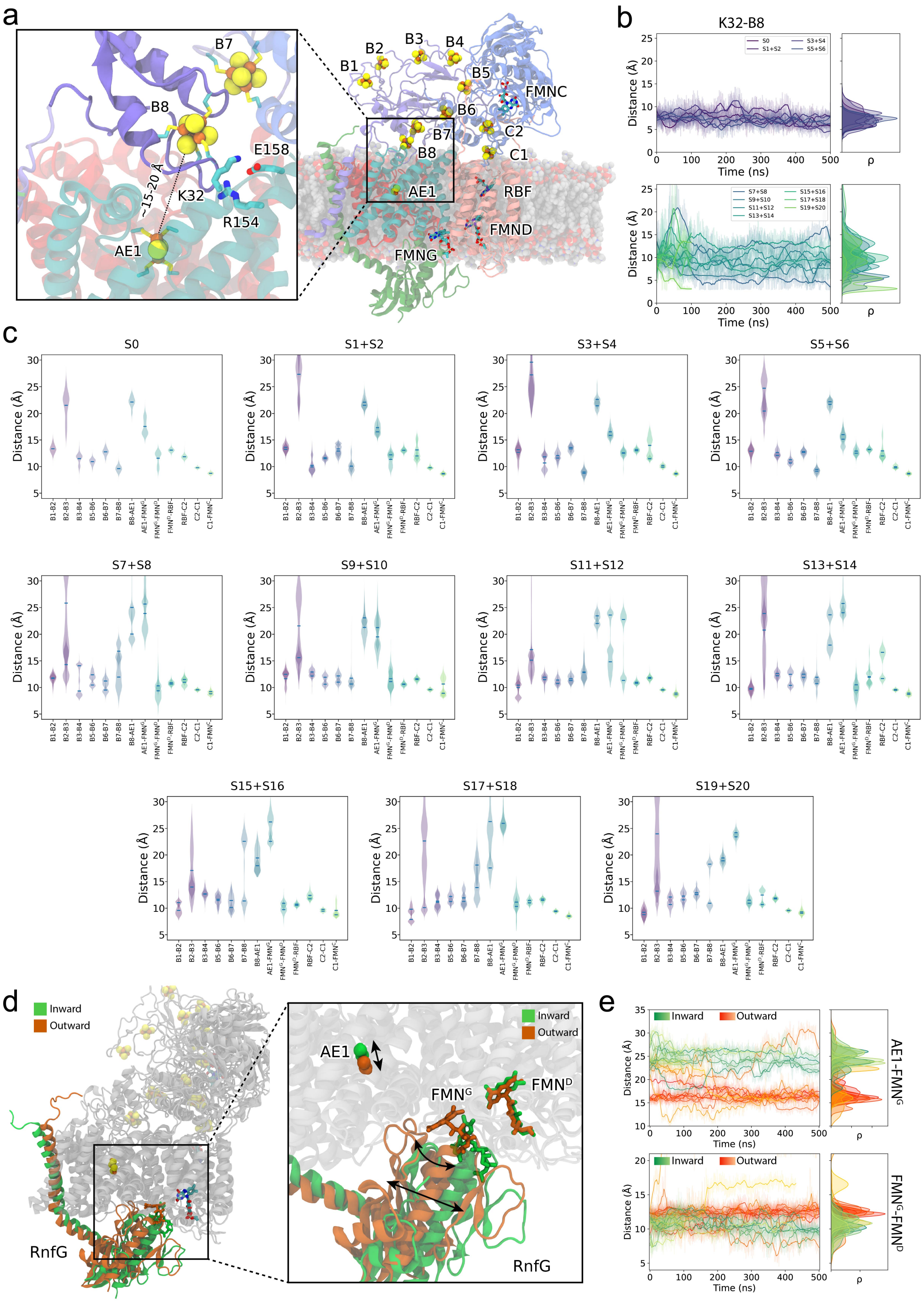

**Extended Data Fig. 9 | Summary of cofactor distances from MD simulations. (a)** Structural overview of the Rnf showing the location of the cofactors after MD equilibration (S13). *Inset:* A snapshot from MD simulation S13 showing the flexible domain of RnfB, containing the B8 cluster, binding close to the membrane subunits of RnfA and RnfE. Two positively charged residues (Lys32$^{RnfA}$, Arg154$^{RnfA}$) are involved in electrostatic interaction with the ISC. **(b)** Distances between Lys32$^{RnfA}$ and B8-cluster of MD simulations starting from NADH- (*top*) or Fd-reduced (*bottom*) structures. See Extended Data Table 6 for the simulation details. **(c)** Violin plots showing all distances between the cofactors. **(d)** Snapshot of RnfG from MD simulations with RnfA/E either in *inward* (green) or in *outward* (orange) conformation. *Inset:* closeup view of the cofactor FMN$^{G}$ showing two distinct conformations. In *outward*-conformation it moves closer to the AE1-center, whereas in *inward*-conformation it moves closer to FMN$^{D}$. **(e)** *Edge-to-edge* distance analysis shown for AE1-FMN$^{G}$ (*top*) and FMN$^{G}$-FMN$^{D}$ (*bottom*) from all MD simulations grouped either into *inward* (green hues) or *outward* (orange hues) conformation.

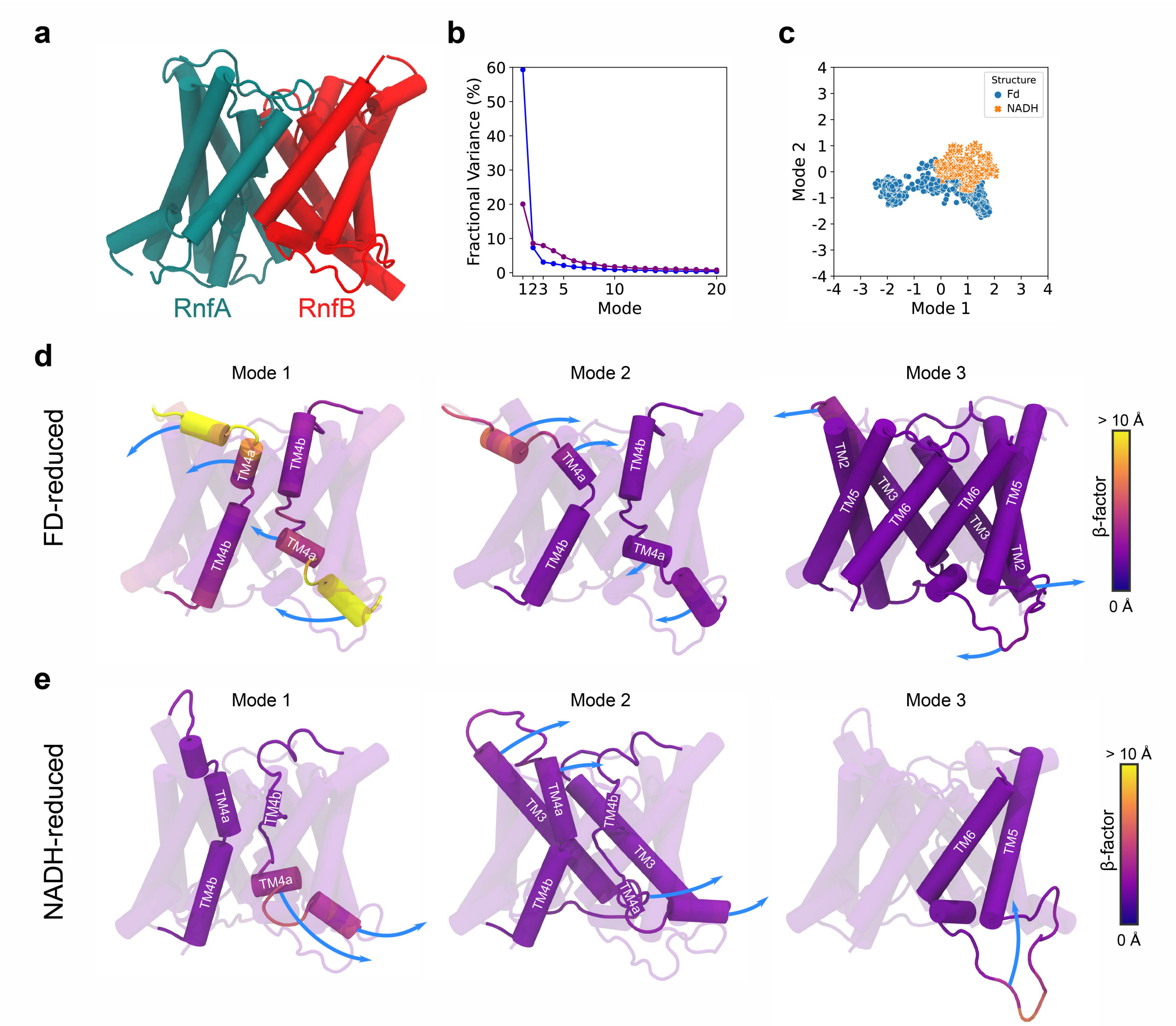

**Extended Data Fig. 10 | Global dynamics of the Rnf complex from principal component analysis (PCA) of the MD simulations. (a)** Overview of subunits used for the PC analysis. RnfA is shown on the left (in *teal*) and RnfB to the right (in *red*). **(b)** Scree plot showing the fractional variance of the obtained normal modes. **(c)** Projection of the MD simulations onto the principal components (PC) - modes 1 and 2. **(d)** The main PC modes of MD simulations based on the Fd-reduced structure with the largest movements of helices TM1 and TM4, showing symmetric (mode 2) or asymmetric (mode 1) motion. Rotational motion of the outer helices (TM2, TM3, TM5) (mode 3). **(e)** The main PC modes from MD simulations of the NADH-reduced structure show overall similar dynamics as in the Fd-reduced simulations, but lower β-factors, indicating a smaller movement. See Extended Data Movies 3 and 4 for the conformational switching and PCA.

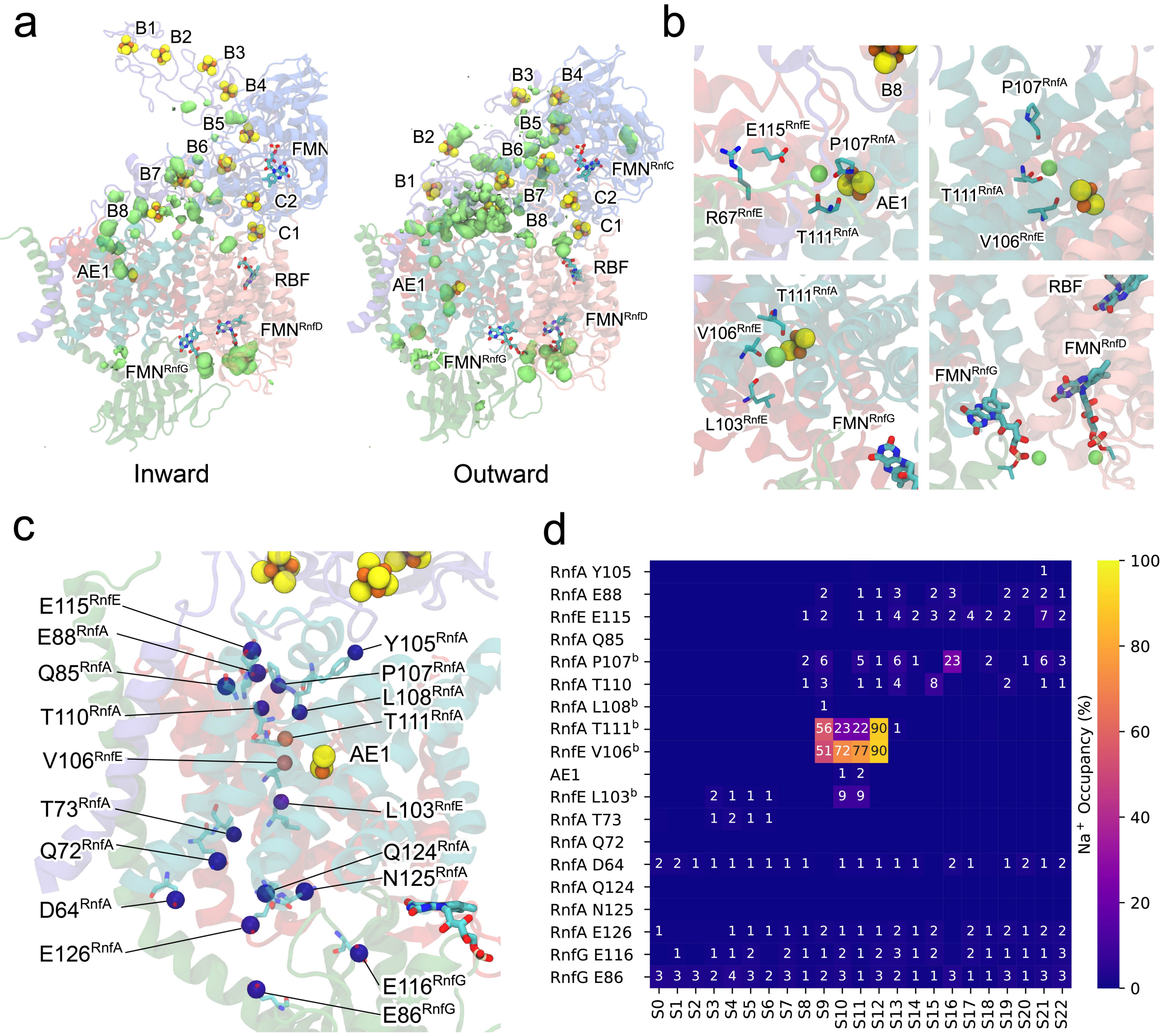

**Extended Data Fig. 11 | Sodium binding in the membrane domain of Rnf. (a)** Analysis of the $Na^+$ occupancy from combined MD simulations of the inward (*left*) and outward (*right*) conformations. The average $Na^+$ occupancies were calculated using the VolMap tool in VMD[45] shown as green-coloured densities. **(b)** Snapshots from MD simulations showing key residues interacting with the $Na^+$ ion. *Top, left*: the cytosolic side, with $Na^+$ ion bound between $E115^{RnfE}$ and $T111^{RnfA}$. *Top, right*: buried binding site in the *inward* conformation, with the $Na^+$ ion bound to the carbonyl backbone of $T111^{RnfA}$ next to the AE1 cluster. *Bottom, left*: the outward conformation, with $Na^+$ ion bound next to the carbonyl backbone of $V106^{RnfE}$ and $L103^{RnfE}$. *Bottom, right:* $Na^+$ ion next to the phosphate groups of $FMN^{RnfG}$ and $FMN^{RnfD}$ on the extracellular side. **(c)** Subunits RnfA/E with potential sodium ion interactions marked in purple spheres. **(d)** $Na^+$ occupancy during MD simulations with the residues shown in panel **c** (see Extended Data Table 6). The occupancies were calculated based on the fraction of frames with a sodium ion within 3 Å of a particular residue (*b* - for only backbone interaction).

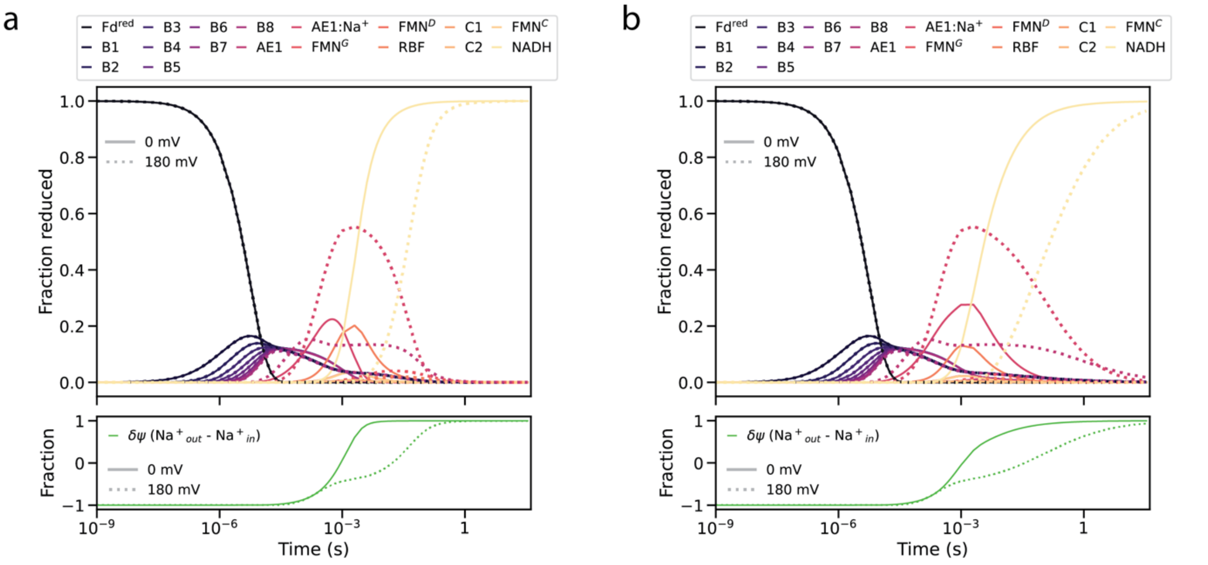

(a) $AE1{:}Na^{+} + FMN^{G}_{ox} \longrightarrow AE1_{ox} + Na^{+}_{out} + FMN^{G}_{ox}$

(b) $AE1{:}Na^{+} + FMN^{G}_{ox} \rightleftarrows AE1_{ox} + Na^{+}_{out} + FMN^{G}_{ox}$

**Extended Data Fig. 12 | Kinetic simulations of the sodium translocation process** with **(a)** an irreversible, and **(b)** a reversible $Na^{+}$ release steps.

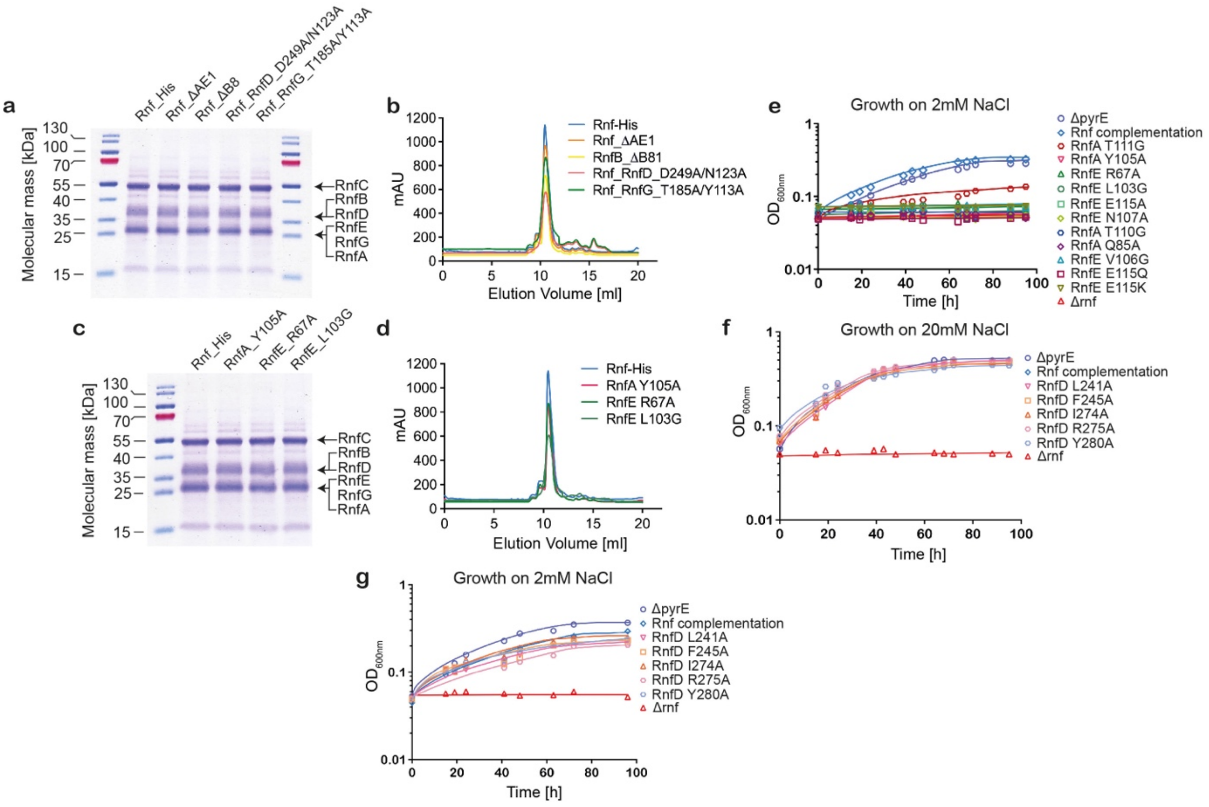

**Extended Data Fig. 13 | Purification and characterisation of Rnf variants from *A. woodii*. (a, b)** 10 µg of Rnf electron transfer pathway variants containing a His-tag purified from *A. woodii* was separated in an SDS-PAGE. Size exclusion chromatography profile of the purified variant Rnf complexes from *A. woodii* on "Superdex 200 Increase™ 10/300". **(c, d)** 10 µg of Rnf variants with alterations in the proposed $Na^+$ binding site containing a His-tag purified from *A. woodii* was separated in an SDS-PAGE. Size exclusion chromatography profile of the purified mutant Rnf complex from *A. woodii* on "Superdex 200 Increase™ 10/300". **(e)** Rnf strain containing mutations of residue involved in $Na^+$ binding and translocation (in RnfAE subunits) did not grow on $H_2$ and $CO_2$ at 2 mM NaCl **(f, g)** Rnf strain containing mutations of residue in RnfD subunits grew as wild type when grown on 20 mM and 2 mM NaCl with $H_2$ and $CO_2$.

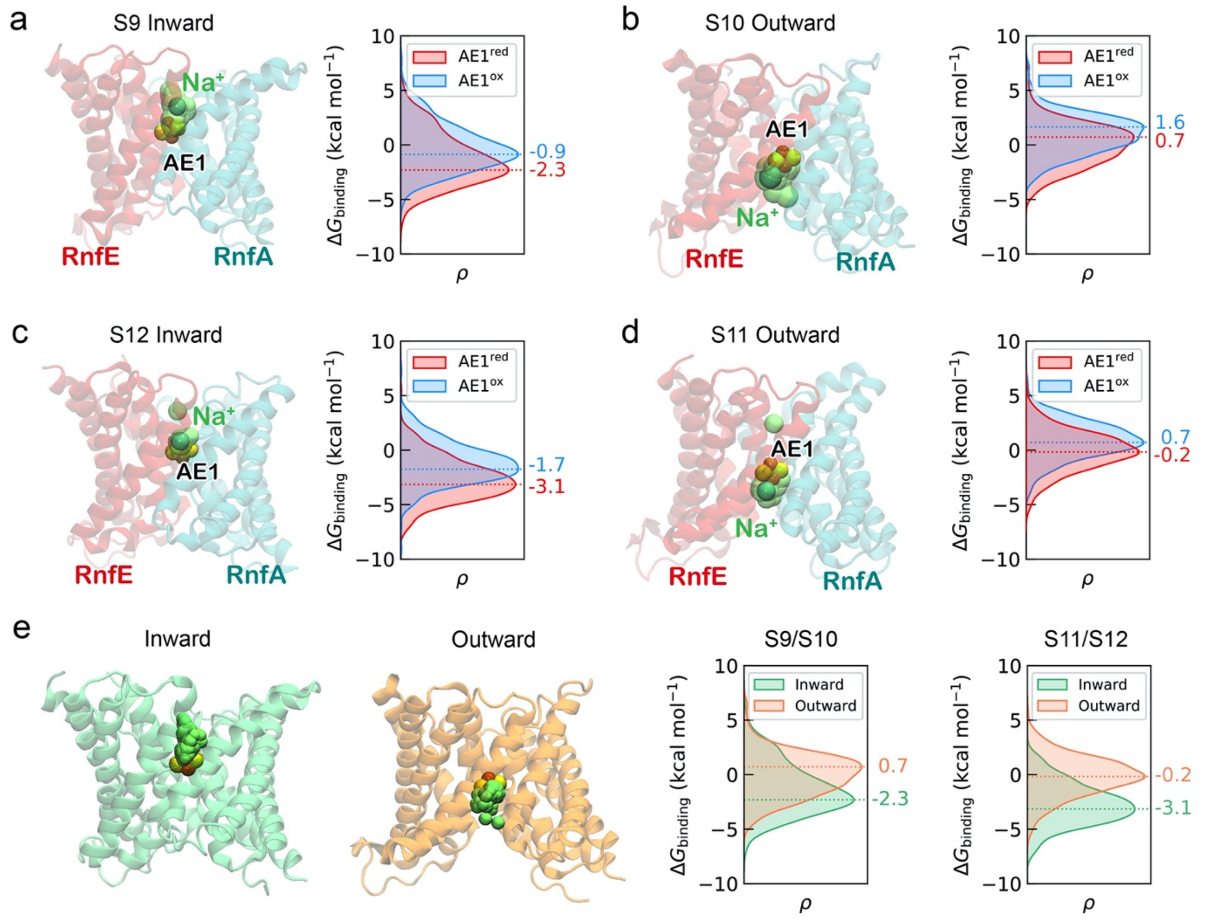

**Extended Data Fig. 14 | Sodium binding to the RnfA/E dimer. (a,b,c,d)** The data shows the binding affinity of $Na^+$ ions in different redox states of the AE1 cluster (red: reduced; blue: oxidised), and in different alternate-access conformations of the RnfA/E dimer. Only snapshots with $Na^+$ bound were considered. The clustered snapshots were generated from MD simulations performed with the reduced AE1 cluster (simulations S9-S12, see Extended Data Table 6), whereas the effect of the AE1 oxidation was modelled by switching the AE1 charges to the corresponding oxidised state, as no spontaneous $Na^+$ binding was observed in MD simulations with an oxidised AE1 cluster. **(e)** Simulations grouped by *inward* (green) and *outward* (orange) conformations, showing the data with reduced AE1 cluster.

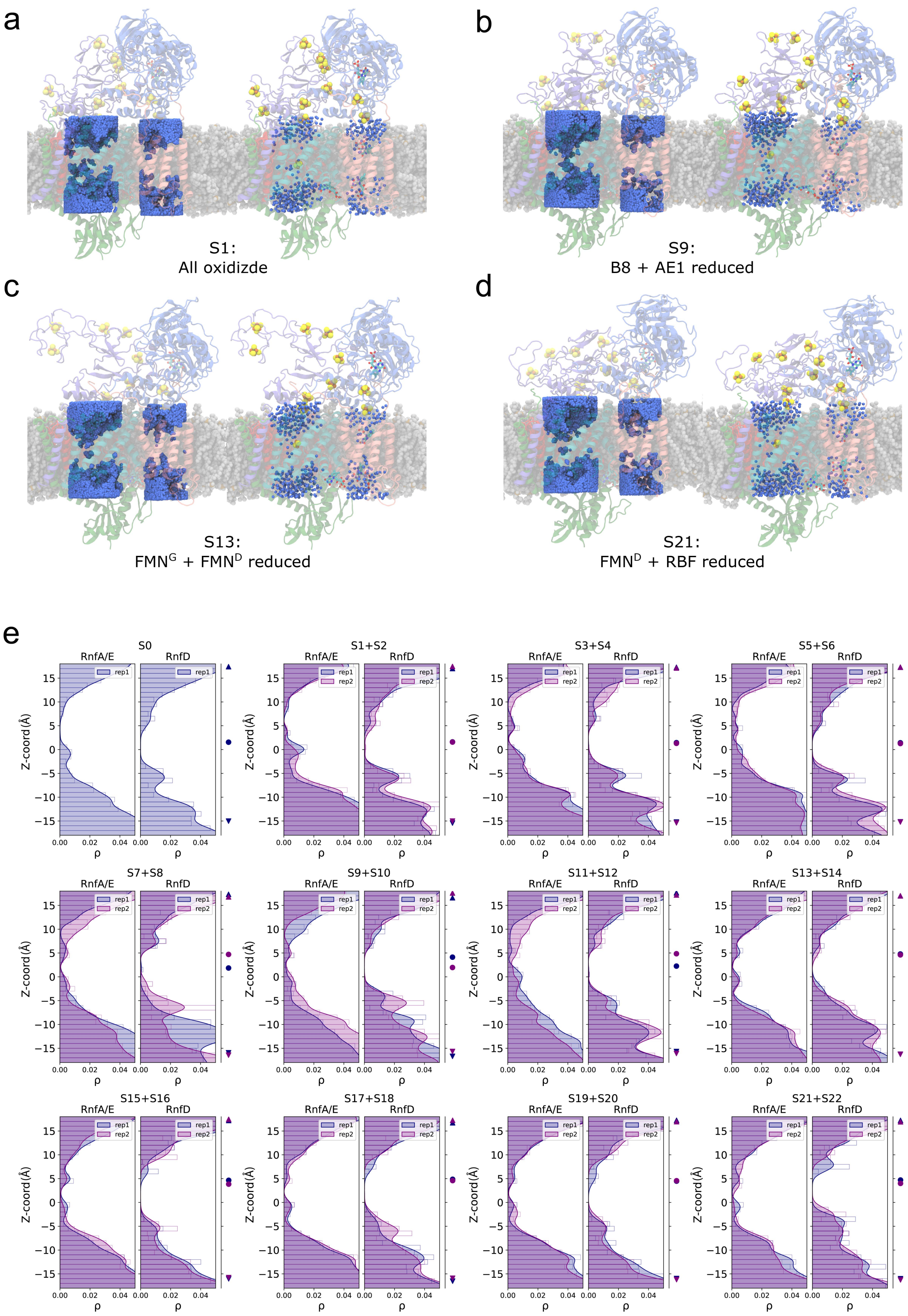

**Extended Data Fig. 15 | Overview of hydration in the membrane subunits of Rnf. (a,b,c,d)** *Left:* Hydration averaged over the MD ensemble, showing water molecules in the subunits RnfA, RnfE, and RnfD. The final 100 ns of each MD simulation with 1 ns/frame are shown. *Right:* The hydration of a final snapshot from the MD simulations. The water selection was defined by a cylinder cantered either to the subunits RnfA/E or RnfD, with the radius of the shape set to $r$ = 14 Å or $r$ = 11 Å, respectively. **(e)** Density of water molecules in RnfA/E (*left*) and RnfD (*right*) projected onto the Z-coordinate (in Å, perpendicular to the membrane plane). The mean position of the upper and lower membrane leaflet, and the AE1 cluster are indicated by triangles and circles, respectively.

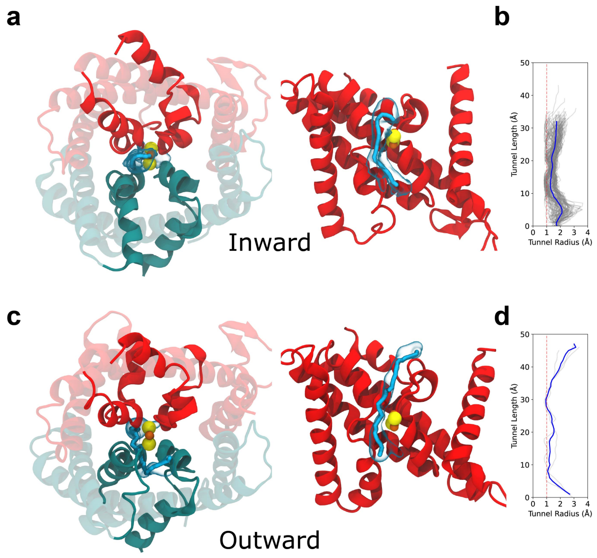

**Extended Data Fig. 16 | Ion channel analysis in RnfA/E. (a, c)** Ion translocation pathways, characterised using tunnel analysis in CAVER[57], showing a continuous pathway from (**a**) the *inward* and **(c)** *outward* sides of RnfA/E. **(b,d)** The tunnel radius of the individual pathways (in grey) and the average tunnel radius (*blue*) in the **(b)** *inward* and (**d**) outward conformations. The analysis was performed on MD simulations S9 and S10 (see Extended Data Table 6).

**Extended Data Movies**

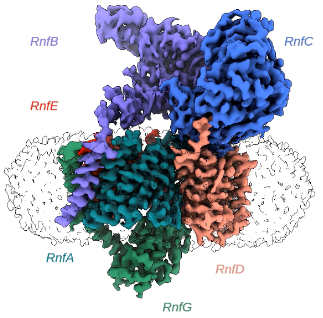

**Movie 1 | Representation of the segmented cryo-EM map of Rnf and its corresponding model. Colour codes used are according to Fig. 1.**

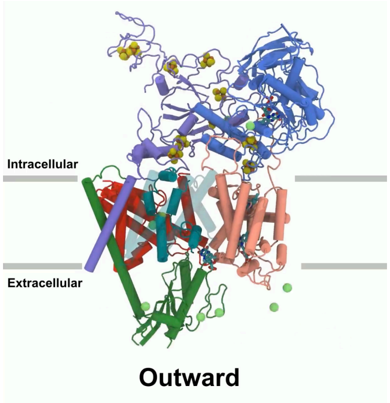

**Movie 2 | Sodium binding from the intracellular and extracellular sides.**

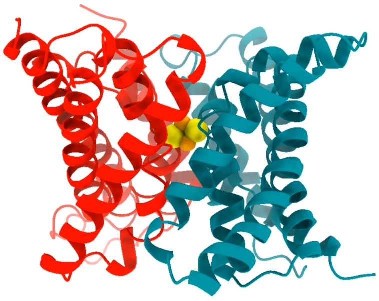

**Movie 3 | Inward/outward transition from MD simulations.**

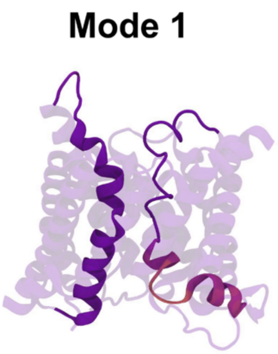

**Movie 4 | Dominant normal modes from MD simulations of the NADH-reduced structures.**

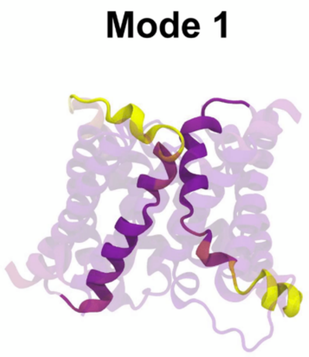

**Movie 5 | Dominant normal modes from MD simulations of the Fd-reduced structures.**

### Extended Data References
